# A novel role for the tumor suppressor gene *ITF2* in lung tumorigenesis and chemotherapy response

**DOI:** 10.1101/517169

**Authors:** Olga Pernía, Ana Sastre-Perona, Carlos Rodriguez-Antolín, Alvaro García-Guede, María Palomares-Bralo, Rocío Rosas, Darío Sanchez-Cabrero, Patricia Cruz, Carmen Rodriguez, MDolores Diestro, Rubén Martín-Arenas, Verónica Pulido, Pilar Santisteban, Javier de Castro, Olga Vera, Inmaculada Ibáñez de Cáceres

## Abstract

Despite often leading to platinum resistance, platinum-based chemotherapy continues to be the standard treatment for many epithelial tumors. In this study we analyze the cytogenetic alterations that arise after cisplatin treatment providing novel insights into the molecular biology and the cellular mechanisms involved in the acquired resistance in these tumor types.

**Methods:** In this study, we used 1 million array-CGH and qRT-PCR methodologies to identify and validate cytogenetic alterations that arise after cisplatin treatment in four lung and ovarian paired cisplatin-sensitive/resistant cell lines. We used whole transcriptome sequencing (RNA-seq), functional transfection assays and gene-pathway activity analysis in our experimental cellular models and in fresh frozen primary NSCLC tumors to identify genes with a potential role in the development of this malignancy. Results were further explored in 55 lung and ovarian primary tumors and control samples and in two extensive in silico databases (TCGA and KMplotter) with 1,926 NSCLC and 1,425 additional epithelial tumors.

**Results:** Long-term cell exposure to platinum induces the frequent deletion of *ITF2* gene. Restoration of *ITF2* expression re-sensitizes tumor cells to platinum and recovers the levels of Wnt/β-catenin transcriptional activity. *ITF2* expression was also frequently downregulated in NSCLC, ovarian and other epithelial tumors, predicting a worse overall survival. We also identified an inverse correlation in expression between *ITF2* and *HOXD9*, revealing that NSCLC patients with lower expression of *HOXD9* have a better overall survival rate that was independent of the tumor histology.

**Conclusion:** We have defined the implication of *ITF2* as a molecular mechanism behind the development of cisplatin resistance probably through the activation of the Wnt-signaling pathway. Our translational data suggest that *ITF2* could be used as a general epithelial tumor platinum-predictive marker and have identified *HOXD9* as a potential prognostic biomarker in NSCLC, a gene which expression is induced by *Wnt* signaling. Furthermore, this data highlights the possible role of *ITF2* and *HOXD9* as a novel therapeutic target for platinum resistant tumors.

## INTRODUCTION

Although platinum-based chemotherapy still plays an important role in the treatment of many solid tumors, the disease progresses to a platinum-resistant state in a high percentage of the diagnosed cases of non-small cell lung cancer (NSCLC) and ovarian cancer ^8, 19^ which are two of the most deadly cancers plaguing our society, the former accounting for more than 80% of primary lung-cancer cases and the latter boasting the highest mortality of the gynecological malignancies worldwide ^14^. Cisplatin (CDDP) is a platinum compound widely used in the treatment of solid tumors. It induces apoptosis in cancer cells by binding to the N7 position of the guanines and crosslinking DNA ^10, 15^. However, CDDP also leads to cytogenetic alterations, such as deletions or amplifications of genes involved in tumor progression, metastasis and drug response ^1^, which contribute to the development of CDDP-resistance ^11, 20, 40^.

In this study, we performed high resolution million feature array-Comparative Genomic Hybridization (aCGH) with four NSCLC and ovarian cancer sensitive/resistant paired cell lines previously reported by our group ^37^ to explore the chromosomic deletions that differ in the resistant subtypes. We found a common deletion that includes the Transcription Factor 4, *TCF4* (hereafter called *ITF2*). *ITF2* is a downstream target gene of the Wnt/β-catenin pathway, that negatively regulates its activity ^18, 35^. Wnt signaling has been identified as one of the key signaling pathways in cancer, and more recently, also to be involved in drug resistance of primary tumors such as colon or ovarian cancer ^6, 26^. However, its role driving platinum resistance in NSCLC has not been defined yet and very little is known about how ITF2 is involved in tumorigenesis. Therefore, further studying of the role of *ITF2* in tumor response to chemotherapy may provide new ways to fight resistance to this popular treatment.

Here we report the frequent downregulation of *ITF2* in NSCLC patients and in cisplatin-resistant cancer cells. Furthermore, we present potential molecular mechanisms, including the Wnt-signaling pathway behind the development of resistance through the action of *ITF2* and affecting the expression of specific genes that might be also used as potential therapeutic targets.

## RESULTS

### ITF2 is frequently downregulated by chromosomal deletion after CDDP cell exposure

We used CDDP sensitive and resistant clones from NSCLC (H23 and H460) and Ovarian cancer (A2780 and OVCAR3) cell lines previously generated by our group to performed CGH arrays and uncover deletions or amplifications that could explain the Cisplatin-resistant phenotype. Our cytogenetic study showed different genomic alterations in the CDDP resistant subtypes, using the sensitive parental cell lines as a reference genome. We found two deleted regions shared by both tumor types in at least three of the four cell lines (H23R, A2780R and OVCAR3R) located on 18q21.2-18q21.31 and 18q21.32, affecting the genes *RAB27B, CCDC68, TCF4, TXNL1, WDR7* and *BOD1P;* and genes *ZNF532, SEC11C, GRP* respectively. We also observed a common deleted region on 2q22.1 that included the gene *LRP1B* in the NSCLC cell lines and an additional common region only shared by the ovarian cancer cell lines on 9q22.33 that included part of the gene *TMOD* (Table S1).

We selected *LRP1B* and *TCF4 (ITF2)* genes that were completely deleted in the same tumor type or in at least three of the four cell lines respectively (Figures 1A and S1A). The deletion of *ITF2* in H23R and A2780R cell lines resulted in a significant loss of *ITF2* expression compared to the sensitive subtypes, validating the results obtained in the arrays CGH (Figure 1B). No changes were observed in OVCAR3 cells for *ITF2* expression or in H23 and H460 cells for *LRPB1* (Figure 1B and S1B).

**Figure 1.**
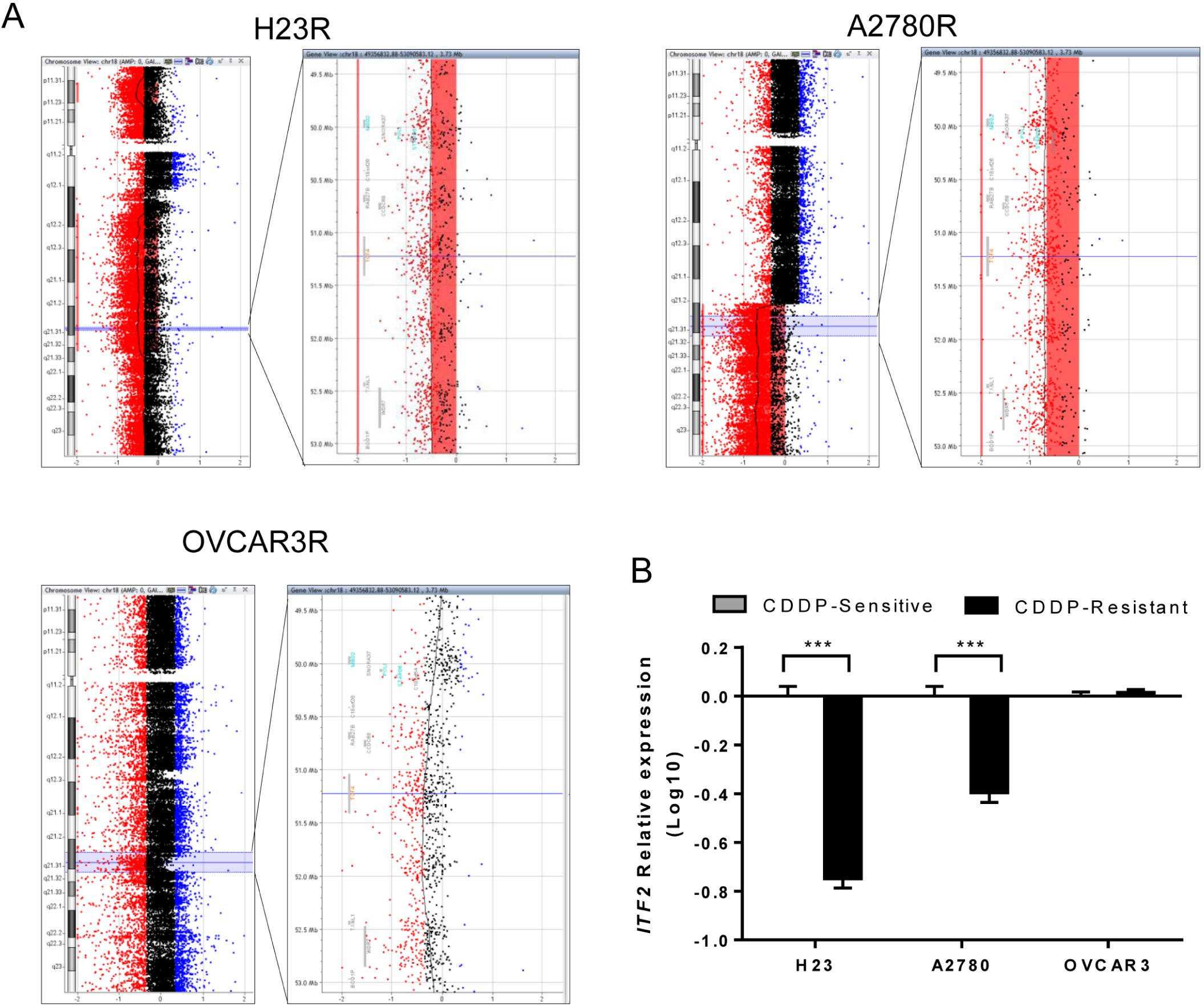
Identification of a common deletion in chromosome 18 in cisplatin-resistant cancer cell lines. (A) Picture extracted from the Agilent Cytogenomics 3.0.1.1 software showing the *ITF2* deletion in chromosome 18 in H23R, A2780R and OVCAR3R cell lines. (B) Relative mRNA expression levels of *ITF2* measured by qRT-PCR. The results show the mean fold induction compared to the sensitive cells. Gene expression was normalized to *GAPDH*. S: sensitive; R: resistant; Data represent the relative expression levels obtained from the combination of two independent experiments measured in triplicate ± SD. *** p < 0.001; (Students T-test).

### Restoration of ITF2 increases the sensitivity to CDDP and decreases β-catenin/TCF transcriptional activity

Due to the association of *ITF2* with the Wnt/ß-catenin/TCF pathway, we studied the transcriptional activity of the Wnt pathway in H23S/R and A2780S/R cells, in which the cisplatin-resistant phenotypes harbor the *ITF2* deletion. We transfected cells with the Wnt reporters (Super8xTop-Fop vectors) and induced the ß-catenin/TCF4 activity either by LiCl treatment, which inhibits GSK3-ß, or by co-transfection with the constitutively stable ß-catenin-S33Y mutant. We observed higher luciferase activity in A2780R cells compared with the parental sensitive ones, indicating an increased transcriptional activity of ß-catenin/TCF in response to both pharmacological (Figure 2A) and functional activation of the pathway (Figure 2B). We also analyzed the RNA-seq normalized FPKM values in paired cell lines, confirming the significant decrease of *ITF2* expression in the resistant phenotype and observing an increase in the expression levels of several downstream effectors genes of this pathway such as *DKK1* (p=0.041), *TCF7L1* (p=0.027) and *TCF7L2* (ns) (Figure 2C), being *DKK1* the most evident in terms of FKPM values. These effects were not observed in H23S/R paired cell lines (Figure 2D).

**Figure 2.**
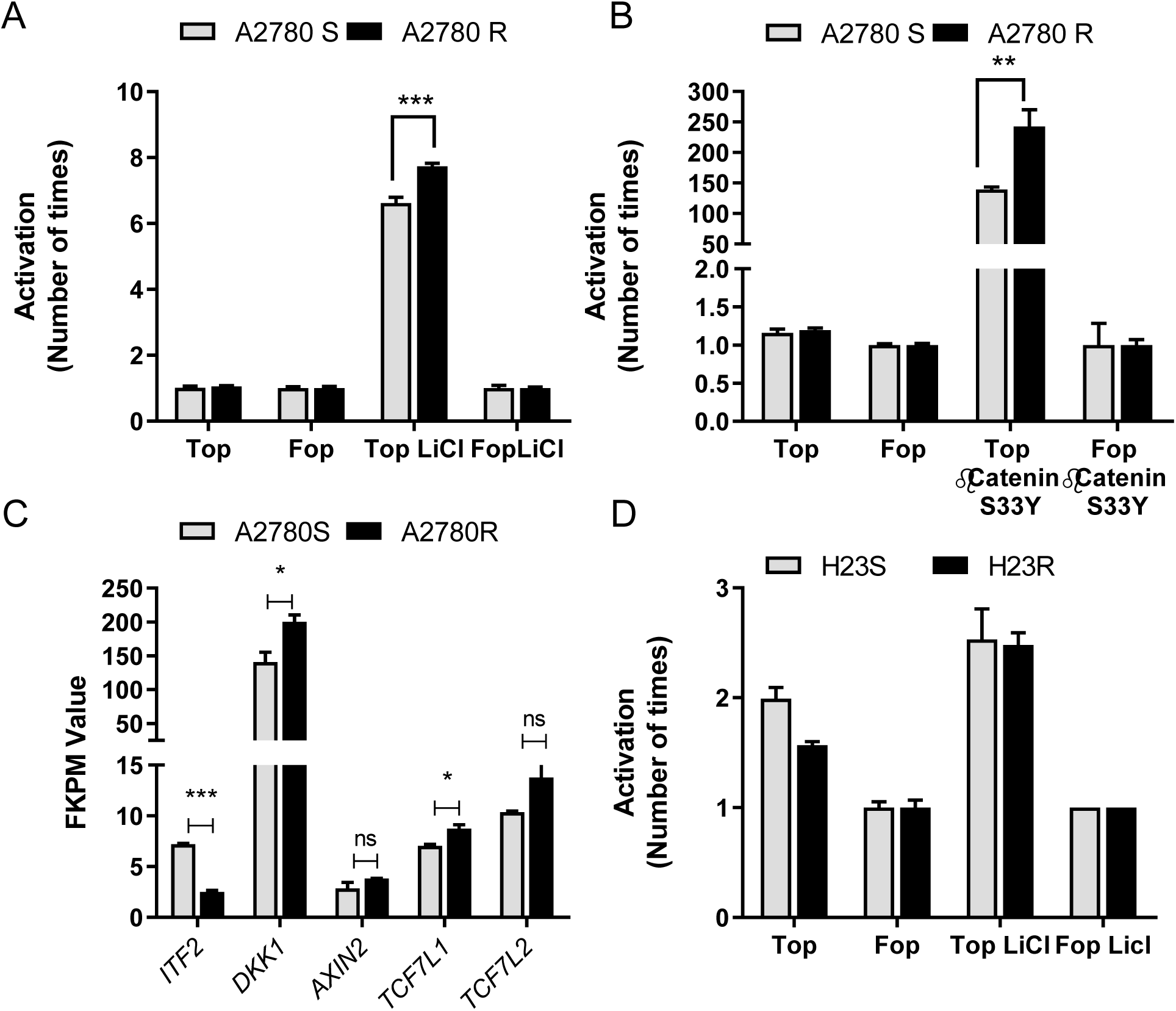
Basal status of the Wnt signaling pathway in A2780 and H23 cell lines. (A-B) Pharmacological activation (A) and functional activation (B) of β-catenin transcriptional activity in A2780 cells. (C) Expression levels of downstream genes regulated by *ITF2* involved in Wnt signaling pathway in A2780S and A2780R cells measured by RNAseq in terms of “Fragments Per Kilobase of transcript per Million mapped reads” or FPKM values (D) Pharmacological activation of β-catenin transcriptional activity in H23 cells. β-catenin transcriptional activity was measure in A2780 and H23 cells after treatment with LiCl (10mM) 24 hours or transfection of bcat-S33Y, transfecting with Super8xTopFlash (Top) or Super8xFopFlash (Fop). The results show the fold induction of the Top/Fop ratio with respect to untreated cells (=1). Values represent the mean of three independent experiments measured by triplicate ± SD. *** p < 0.001; ** p < 0.01, *p<0.05 (Students T-test) ns: non significant.

In order to test the role of *ITF2* in cisplatin resistance through the Wnt pathway, we transiently overexpressed *ITF2* cDNA in A2780 cells, as our previous results confirmed this as a reliable cellular model to evaluate changes in the transcriptional activity of the Wnt pathway. First, we observed that the overexpression of *ITF2* in A2780R resulted in a significant increase in sensitivity to cisplatin from the dose of 0.5ug/ml (p<0.01), showing an intermediate phenotype between the resistant and sensitive subtypes (Figure 3A). In addition, *ITF2* restoration induced a dramatic decrease in cell viability (p<0.05) 24 hours after transfection compared with the parental resistant cells transfected with the empty vector (Figure 3B). *ITF2* overexpression at 24 and 72 hours after transfection was confirmed by qRT-PCR (Figure 3C). Moreover, *ITF2* overexpression recovered the levels of ß-catenin/TCF transcriptional activity observed in sensitive cells (Figure 3D). We also determined *DKK1* expression levels after *ITF2* overexpression as it was the downstream effector gene showing the highest increased expression in the resistant A2780R cells. In fact, its expression was restored in part at 24h (p<0.05) after *ITF2* overexpression (R-ITF2). Decreased levels of DKK1 were also observed at 72h in both, semi-quantitative and quantitative assays, without associated significance, probably due to the in transient transfection (Figure 3E and 3F).

**Figure 3.**
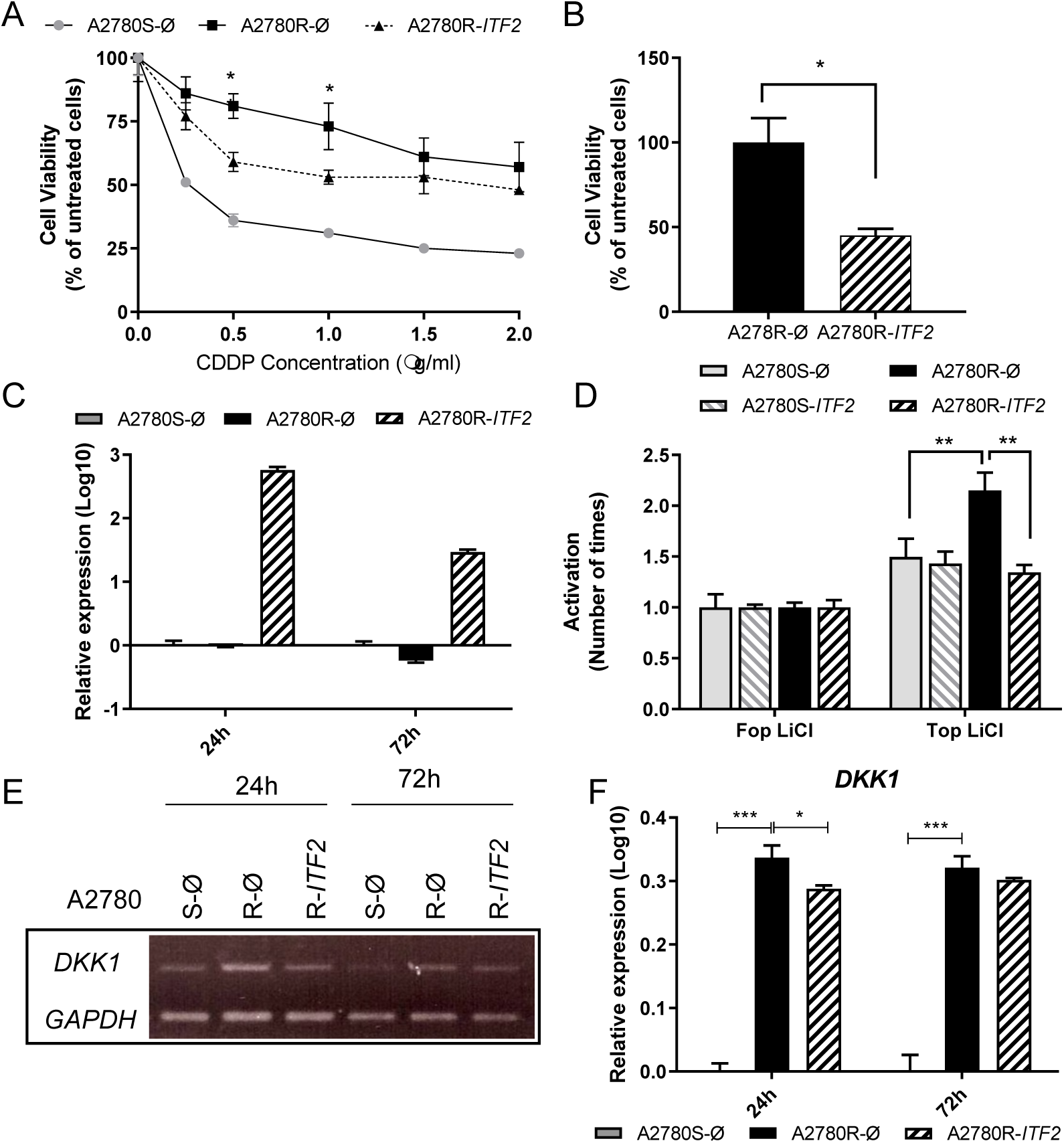
Effect of *ITF2* on cisplatin resistance, cell viability and Wnt pathway. (A) Viability curves of A2780 cell lines transfected with pCMV6 (S-Ø and R-Ø) and with the overexpression vector (R-ITF2). Each experimental group was exposed to six different CDDP concentrations for 48h. Data were normalized to each untreated control, set to 100%. The data represent the mean ± SD of at least three independent experiments performed in quadruplicate at each drug concentration for each cell line analyzed. (B) Viability of A2780 cell lines transfected with pCMV6 (R-Ø) and with the overexpression vectors (R-ITF2). (C) Relative expression levels of *ITF2* measured by quantitative RT-PCR, represented in Log10 scale; In each experimental group, the sensitive cell line transfected with pCMV6 plasmid was used as a calibrator. Each bar represents the combined relative expression of two independent experiments measured in triplicate. (D) β-catenin transcriptional activity was measured in A2780 cells after *ITF2* overexpression and treatment with LiCl (10mM) for 24 hours, transfecting with Super8xTopFlash (Top) or Super8xFopFlash (Fop). The results show the fold induction of the Top/Fop ratio with respect to untreated cells (=1). Values represent the mean of three independent experiments measured by triplicate ± SD. (E) Expression analysis of the downstream gene DKK1 regulated by *ITF2* in A2780 cell line transfected with pCMV6 (S-Ø and R-Ø) and with the overexpression vector (R-ITF2) for 24 and 72 hours. Representative images of *DKK1* and *GAPDH* measured by RT-PCR. (F), Expression levels of *DKK1* measured by qRT-PCR. Each assay was performed at least three times to confirm the results. *** p < 0.001; * p < 0.05 (Students T-test).

### The expression of ITF2 is frequently downregulated in NSCLC, ovarian and other epithelial tumors

To validate our *in vitro* results, we determined the clinical implication of *ITF2* and *DKK1* expression in NSCLC and ovarian cancer patients. The relative expression of both genes was measured in two cohorts of fresh frozen tumor samples (T) and adjacent tissue (ATT) from NSCLC (Table 1) and ovarian cancer patients (Table S2).

**Table 1.** Clinicopathological and experimental data obtained from patients with NSCLC from La Paz University Hospital. Note: OS, Overall Survival; PFS, Progression Free Survival; CDDP: cisplatin; CBDCA: carboplatin; NA: not available.

| Patient | Histology | Sex | Stage | Chemotherapy | Relapse | Status | OS (days) | PFS (days) | TCF4<br>(2<sup>-ΔCt</sup>) | DKK1<br>(2<sup>-ΔCt</sup>) | HOXD9<br>(2<sup>-ΔCt</sup>) |
| --- | --- | --- | --- | --- | --- | --- | --- | --- | --- | --- | --- |
| Pat1.T | Adenocarcinoma | Female | IA | No | Yes | Alive | 2220 | 1490 | 0,03364 | 0,00089 | 0,00008 |
| Pat2.T | Adenocarcinoma | Male | NA | No | Yes | Alive | 2352 | 1860 | 0,01041 | 0,00003 | 0,04277 |
| Pat3.T | Epidermoid | Male | IB | No | Yes | Exitus | 1022 | 825 | 0,01012 | 0,00075 | 0,00001 |
| Pat4.T | Adenocarcinoma | Male | IB | No | No | Exitus | 3 | 3 | 0,00286 | 0,00006 | 0,00041 |
| Pat5.T | Adenocarcinoma | Male | NA | No | No | Exitus | 626 | 626 | 0,00365 | 0,00005 | 0,00503 |
| Pat6.T | Large cell | Male | IIB | No | No | Exitus | 62 | 62 | 0,01584 | 0,00119 | 0,00074 |
| Pat7.T | Adenocarcinoma | Male | IIIA | Otro | Yes | Exitus | 228 | 138 | 0,01278 | 0,00150 | 0,00078 |
| Pat8.T | Epidermoid | Female | IIIB | CDDP + Others | No | Exitus | 109 | 109 | 0,00675 | 0,00026 | 0,00126 |
| Pat9.T | Adenocarcinoma | Female | IIA | CDDP + Others | Yes | Alive | 2260 | 2260 | 0,01070 | 0,01826 | 0,00097 |
| Pat10.T | Epidermoid | Male | IB | No | No | Alive | 1853 | 1853 | 0,01626 | 0,00162 | 0,00027 |
| Pat11.T | Adenocarcinoma | Male | IA | No | No | Exitus | 216 | 216 | 0,00456 | 0,00003 | 0,00070 |
| Pat12.T | Adenocarcinoma | Female | IIIA | CBDCA + Others | Yes | Alive | 2192 | 2192 | 0,00580 | 0,00004 | 0,00146 |
| Pat13.T | Epidermoid | Male | IB | CDDP + Others | No | Alive | 2341 | 2341 | 0,02466 | 0,00090 | 0,00013 |
| Pat14.T | Epidermoid | Male | IIA | No | ND | Exitus | 289 | 289 | 0,01179 | 0,00206 | 0,00061 |
| Pat15.T | Epidermoid | Male | IIA | No | ND | ND | 109 | 109 | 0,01833 | 0,01370 | 0,00420 |
| Pat16.T | Adenocarcinoma | Female | IIIA | CDDP + Others | ND | Alive | 2228 | 2228 | 0,00907 | 0,00005 | 0,00191 |
| Pat17.T | Adenocarcinoma | Male | IIB | Otro | Yes | Exitus | 888 | 443 | 0,00704 | 0,00014 | 0,00002 |
| Pat18.T | Epidermoid | Male | IIB | CBDCA + Others | No | Exitus | 259 | 259 | 0,00563 | 0,00001 |  |
| Pat19.T | Adenocarcinoma | Female | IB | CDDP + Others | Yes | Exitus | 936 | 428 | 0,00738 | 0,00137 | 0,00014 |
| Pat20.T | Epidermoid | Male | IIB | CDDP + Others | Yes | Exitus | 1224 | 637 | 0,00346 | 0,00089 | 0,00264 |
| Pat21.T | Adenocarcinoma | Male | IIIA | CDDP + Others | No | ND | 421 | 421 | 0,00792 | 0,00279 | 0,00003 |
| Pat22.T | Adenocarcinoma | Female | IIB | CDDP + Others | ND | ND | 184 | 184 | 0,01319 | 0,00101 | 0,00007 |
| Pat23.T | Epidermoid | Male | IIIA | CDDP + Others | ND | ND | 542 | 542 | 0,03221 | 0,00214 | 0,00120 |
| Pat24.T | Adenocarcinoma | Male | IIA | CBDCA + Others | No | Alive | 1491 | 1491 | 0,03680 | 0,00614 | 0,00077 |
| Pat25.T | Adenocarcinoma | Male | IIA | CDDP + Others | No | Alive | 1496 | 1496 |  |  | 0,00008 |

We observed that *ITF2* expression is frequently downregulated in NSCLC and ovarian tumor samples (Figure 4) validating our *in vitro* data. Fifteen out of 25 tumor samples of NSCLC patients had lower expression of *ITF2* compared to the normal lungs mean (NLM) (Figure 4A). Furthermore, as reported in our experimental data, we observed the opposite expression profile between *ITF2* and *DKK1* in 60% of NSCLC samples. However, this situation was found only in approximately 10% (1 out of 9) of the ovarian cancer samples (Figure 4B). We did not observe differences between ATT and normal lung samples (LC) for *ITF2* (p=0.177) and *DKK1* (p=0.693) in the NSCLC cohort (Figure 4A).

**Figure 4.**
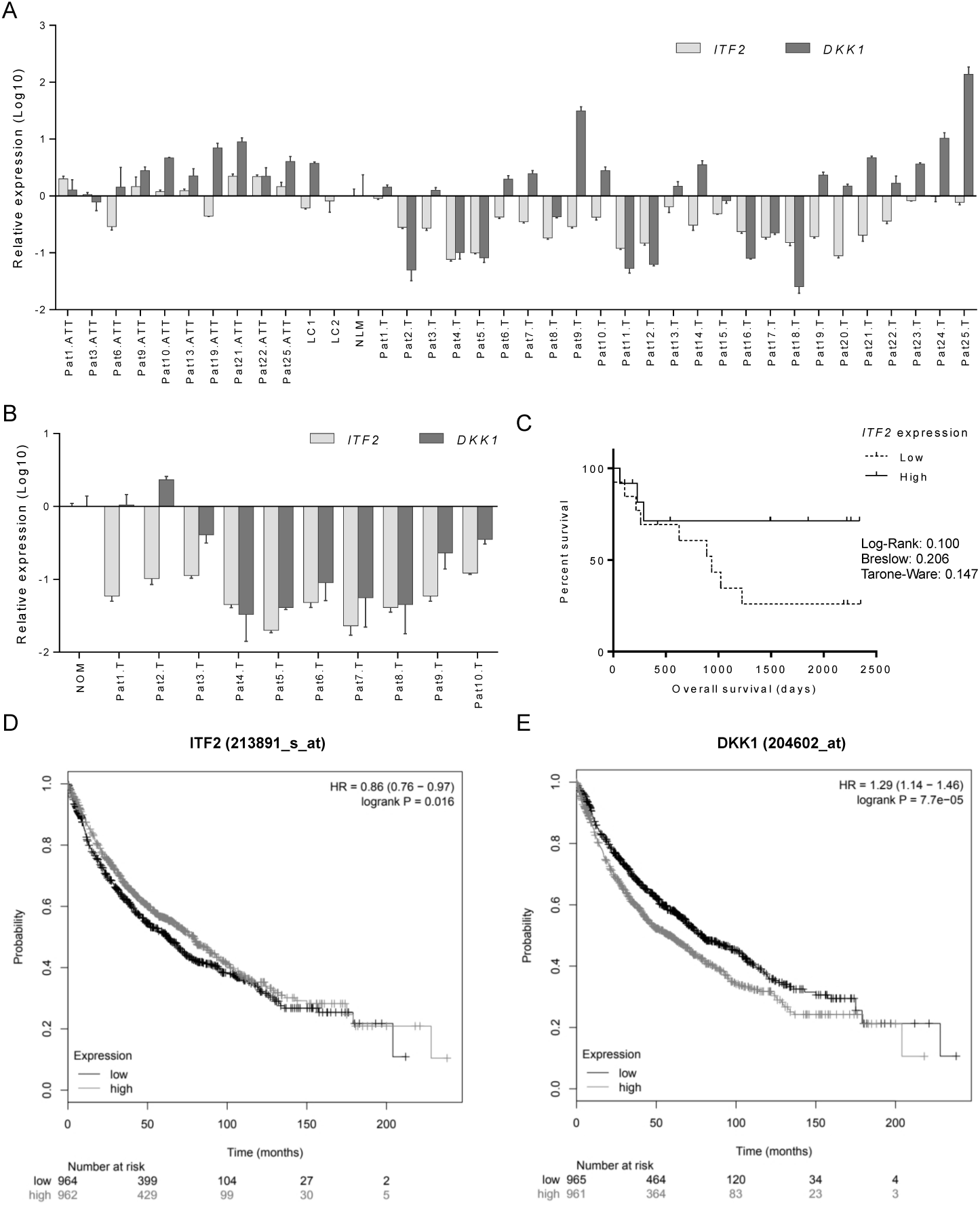
Expression profile of *ITF2* and *DKK1* in patients with NSCLC and ovarian cancer. (A-B) Assessment of *ITF2* and *DKK1* expression levels measured by qRT-PCR in 55 fresh samples from non-tumor samples and two cohort NSCLC (A) and ovarian cancer patients (B). For all the analyses, data represents expression levels in 2-ΔΔCt using the mean of normal lungs (NLM) or ovarian (NOM) as calibrator. (C) Survival analysis in 25 NSCLC samples according to the mean of *ITF2* expression. LogRank, Breslow and Tarone-Ware test were used for comparisons and p<.05 was considered as a significant change in OS. NSCLC: non-small cell lung cancer; ATT: adjacent tumor tissue; T: tumor; LC1/LC2: Lung Control; NLM: Normal Lung Mean; NOM: Normal Ovarian Mean. (D-E) Survival analysis in 1 926 patients from the Kaplan Meier online tool^9^, for ITF2 (D) and DKK1 (E) in terms of overall survival.

The Kaplan Meier curves, analyzing the overall survival (OS) according to the median of *ITF2* and *DKK1* expressions, showed that only *ITF2* high expression levels had a trend towards a better overall survival in NSCLC patients with high *ITF2* expression levels after 250 days of following up (p=0.1) (Figure 4C and Figure S2A). However the statistical significance of both genes in predicting overall survival, was confirmed analyzing an extended cohort of 1,926 lung cancer patients by using the Kaplan Meier plotter online tool, revealing that those patients with high expression of *ITF2* (Figure 4D) and low expression of *DKK1* (Figure 4E) had a significantly better overall survival rate (p=0.016 and p<0.001, respectively). This was validated using the TCGA data set of 487 lung carcinoma (Figure S2B). In addition, *ITF2* was found deleted or downregulated in other epithelial tumors, like Esophageal adenocarcinoma (186 patients) or Head and Neck SCC (522 patients), where *ITF2* loss also predicted a worse overall survival (Figure S2C and D).

### Identification of candidate genes involved in the Wnt signaling pathway through the analysis of RNA-seq in NSCLC patients

To further explore the role of the Wnt-signaling pathway in lung cancer tumorigenesis, we performed a whole transcriptome analysis performing RNA-seq on 14 samples including nine NSCLC samples, six of them with an inverse expression profile between *ITF2* and *DKK1* (Pat3, Pat6, Pat10, Pat22, Pat25 and Pat26) and three with the same expression profile (Pat8, Pat16 and Pat18) (Table S3). Three ATT (Pat9, Pat21 and Pat25) and two normal lung samples (LC1 and LC2), were used as controls for comparisons (Figure S3). Because *ITF2* was mainly downregulated in NSCLC patients, while *DKK1* showed a more heterogeneous expression pattern, we considered *DKK1* as the best parameter to decide the bioinformatics analysis of the RNA-seq. The bioinformatic contrasts focused in three differential gene expression analysis: contrast A, tumor versus control; contrast B, comparison between tumors with high and low expression of *DKK1*; and contrast C, comparison between tumor samples with high expression of *DKK1* and controls (Figure 5). We selected the genes that showed significant expression differences (FDR<0.05) in at least two of the three contrasts, prioritizing contrast B (Table S4). The bioinformatics analysis also focused on all annotated genes related to the Wnt-pathway.

**Figure 5.**
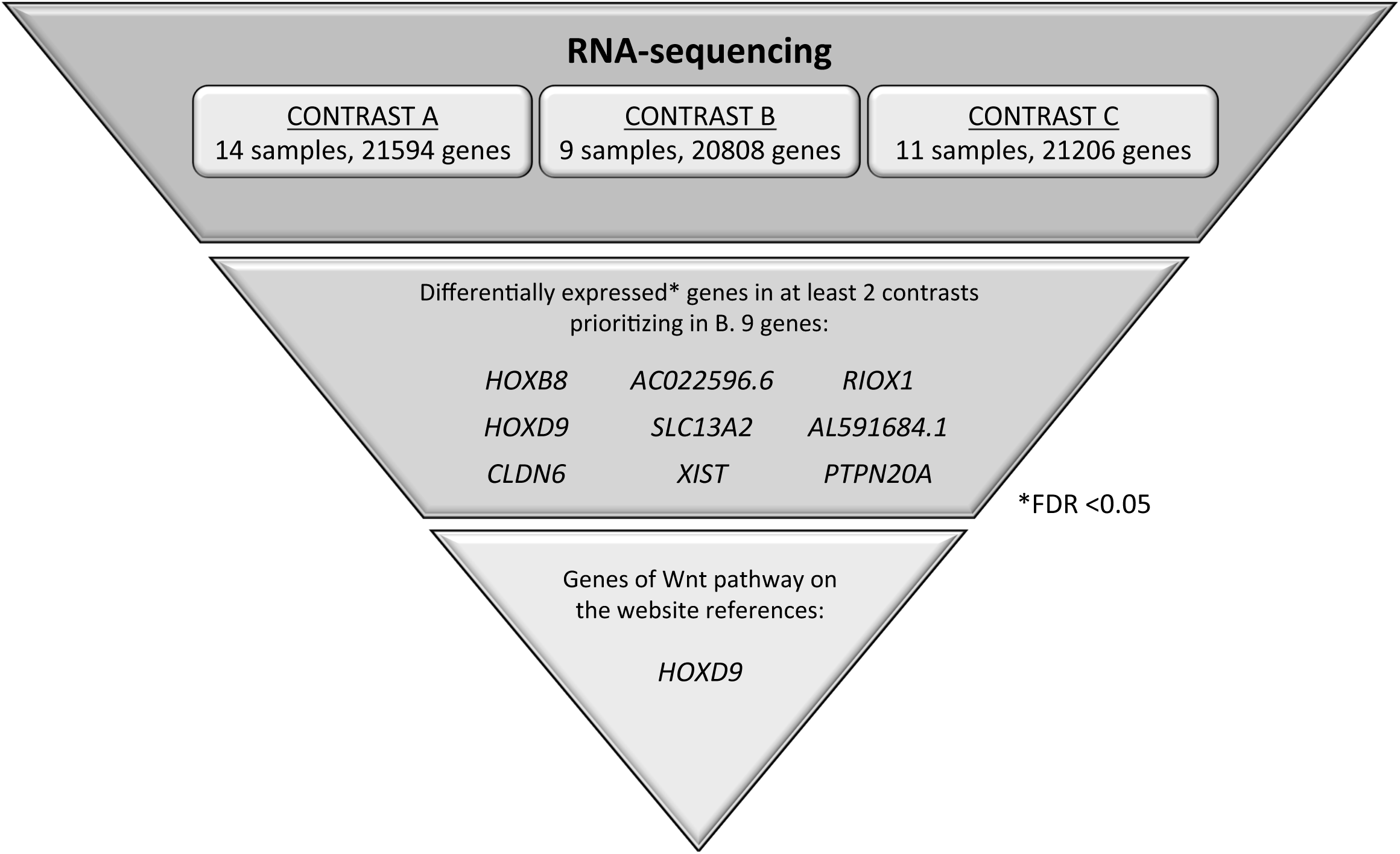
Selected genes potentially involved in the Wnt signaling pathway. **Genes** were identified through a global transcriptomic analysis of 14 NSCLC patient samples combining the information with all annotated genes related to the Wnt-pathway in gene sets of the Molecular Signatures Database (MSigDB, Broad Institute)

We analyzed the expression of nine candidates, but only four of them were confirmed by qPCR-PCR, three coding genes, *HOXD9, RIOX1* and *CLDN6* and one long non-coding RNA, *XIST*. An accurate correlation with the RNA-sequencing data was found for *HOXD9, CLDN6* and *XIST* genes (r=0.83, r=0.97 and r=0.97, respectively) (Figure S4), while for *RIOX1* the correlation coefficient was less marked, probably because of the sample size (r=0.58) (Figure S4). From all four candidates, only *HOXD9* expression showed correlation with *ITF2* expression, (Pearson=-0.24) (Figure 6A). In order to gain insight into the role of *ITF2* regulating the expression of *HOXD9* in NSCLC, we overexpressed *ITF2* in H23 resistant lung cancer cells. Transfection efficiency was confirmed by qRT-PCR at 24 and 72 hours after transfection (Figure 6B). As expected from the primary tumors results, the overexpression of *ITF2* induced a significant decrease of *HOXD9* (p=0.017) (Figure 6C). Having identified *HOXD9* as potential Wnt pathway candidate genes regulated by *ITF2*, we studied its clinical translational application in the cohort of NSCLC patients used for the RNA-seq analysis and in the public databases TCGA and KMplotter.

**Figure 6.**
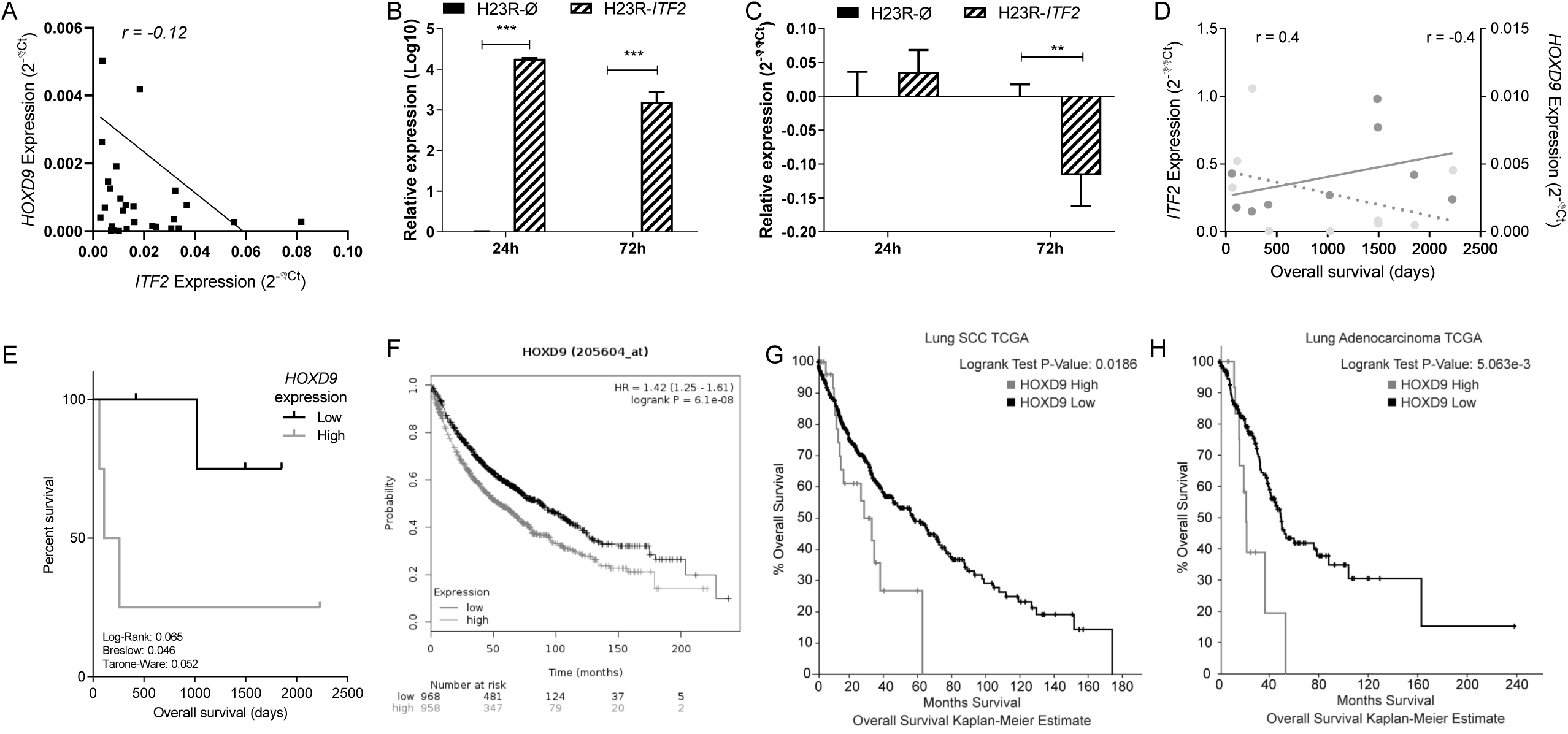
Analysis and clinical significance of *ITF2* and *HOXD9* in NSCLC samples. (A) Correlation between *ITF2* expression and *HOXD9* in tumor and non-tumor samples from the complete cohort of 25 NSCLC patients. Pearson coefficient was used for linear correlation of the quantitative variables. (B-C) Effect of *ITF2* overexpression on *HOXD9* levels (B) Validation of the transfection efficiency of *ITF2* at mRNA levels. Relative expression levels of *ITF2* measured by qRT-PCR, in the cell line H23R, at 24 and 72 hours after transfection represented in Log10 the 2-ΔΔCt. (C) Relative expression levels of *HOXD9* measured by quantitative RT-PCR after *ITF2* overexpression. For both (B) and (C) the resistant cell line transfected with pCMV6 plasmid was used as a calibrator (R-Ø). H23R cells were also transfected with ITF2 cDNA (R-ITF2). Each bar represents the combined relative expression of two independent experiments measured in triplicate. *** p < 0.001; ** p < 0.01 (Students T-test). (D) Correlation between *ITF2* and *HOXD9* expression levels with the overall survival analyzed in NSCLC patients selected form the RNA-seq analysis. Data represents the quantitative expression levels of the two genes measured by qRT-PCR and represented as 2-ΔΔCt for ITF2 (referred to the NLM) and 2-ΔCt for *HOXD9*. (E) Survival analysis in NSCLC samples according to the mean of *HOXD9*. LogRank, Breslow and Tarone-Ware test were used for comparisons and p < 0.05 was considered as a significant change in OS. (F-H) Survival analysis in 1,926 patients from the Kaplan Meier online tool (F) and the TCGA data sets of lung Adenocarcinoma (G, High=23; No Change=203) and SCC (H, High=28; No Change=441) for *HOXD9*.

A negative correlation for *HOXD9* was found in terms of patients’ overall survival (Figure 6D). In addition, the survival analysis performed after stratifying patients according to the median of *HOXD9* gene expression showed that patients with lower expression of *HOXD9* present a significantly better overall survival rate (p=0.046) (Figure 6E). These results were confirmed when analyzing the expression levels of 1,926 lung cancer patients by using the Kaplan Meier plotter online tool and the TCGA data sets, revealing that those patients with lower expression of *HOXD9* had a significantly better overall survival rate that was independently associated with the tumor histology (p<0.001) (Figure 6 F-H).

## DISCUSSION

Wnt signaling has recently been reported to be involved in driving platinum resistance of several tumor types ^6, 41^. However, the molecular mechanisms implicated are not clear, especially in NSCLC. In the present work, we have studied the involvement of the Wnt signaling pathway in tumorigenesis through a combined experimental approach by using both CGH arrays and RNA-sequencing. We have found that long term exposure to platinum induces a frequent deletion of *ITF2* that is involved, at least in part, in the activation of the Wnt pathway.

First, we identified a common deletion in H23, OVCAR3 and A2780 cells, induced by cisplatin treatment in chromosome 18, including two completely deleted genes, *LRP1B* and *ITF2.* A similar deletion was identified in a previous study analysing cisplatin response in ovarian cancer samples, which supports our results ^3^. However, there is another study analyzing the cytogenetic alterations of CDDP-resistant A2780R cells, which shows different genomic alterations. This was probably due to the specificity of the CGH-array used and the experimental design that included less representative number of probes and was performed in only one cell line ^20^. We were not able to validate the *LRP1B* expression changes by using an alternative technique, an issue that has been previously reported ^17^. In our case, it could be due to the mosaicism observed in this region that occurs in less than 20% of the resistant cells. The level of mosaicism that can be detected is dependent on the sensitivity and spatial resolution of the clones and rearrangements present ^7^. Nevertheless, *LRP1B* could still play a role in tumor progression as several studies link its downregulation through deletion and carcinogenesis ^4, 22, 28^. *ITF2* expression changes were confirmed in H23R and A2780R but not in OVCAR3R cells, also probably due to the level of mosaicism (36%) observed in these cells. Our results indicate that low levels of mosaicism would make the validations of expression changes by another quantitative technique difficult, probably because the alterations at expression levels are not significant enough to be detected.

The fact that *ITF2* is deleted and downregulated after platinum treatment provides us with new insight regarding its importance in resistance to platinum chemotherapy in lung and ovarian cancer. In line with our results a previous study, using targeted sequencing in a PDX-based modeling of breast cancer chemoresistance, identified a genomic variant of ITF2 that depicted a link between its altered expression and breast cancer chemoresistance, although no detailed mechanism was provided to connect ITF2 function to chemoresistance ^33^.

ITF2 is a transcription factor belonging to the basic Helix Loop Helix (bHLH) family, which can act as a transcriptional activator or repressor ^25, 34^ but much about its regulation remains unknown. It is important to distinguish *ITF2*, whose official name is TCF4, from the T-Cell Factor 4 (TCF7L2), also known as *TCF4*, which is the ß-catenin transcriptional partner ^27^. In fact, *ITF2* expression is induced by the ß-catenin/TCF complex, but at the same time, it acts as a repressor of this complex by interfering with the binding of B-catenin to TCF4. This causes a decrease in the expression of Wnt target genes, leading to the repression of cell proliferation ^35^. Consistent with these studies, we have observed that the resistant A2780 cells have increased activity of the ß-catenin/TCF transcription, which is concomitant with the increased expression of the downstream effector gene DKK1, probably due to the absence of ITF2. In contrast, we did not observe differences in H23 cells, which could be explained by previous observations ^2^ confirming that H23 has a high basal activity in the Wnt signaling pathway and therefore exogenous activation may not show a difference. Moreover, we have observed that the overexpression of *ITF2* in A2780R cells leads to a decrease in cell viability, rescuing the sensitive phenotype maybe through the inhibition of the excessive proliferation and the activity levels of the B-catenin/TCF transcription. In fact, our results indicate that resistant cells respond better to the activation of the Wnt pathway, an effect that is restored after the re-expression of *ITF2*. Therefore, ITF2 plays an important role in the resistant to cisplatin probably through the regulation of Wnt signaling pathway

Our translational analysis based on the expression levels of *ITF2* and *DKK1* genes in two different cohorts of patients was aimed to elucidate the role of this pathway in tumor progression and chemotherapy response. *ITF2* expression was frequently downregulated in NSCLC and ovarian tumor samples, validating our *in vitro* data. The expression levels of *DKK1*, however, showed a more heterogeneous pattern in the NSCLC tumor samples, while no differences were observed in the ovarian tumors, suggesting an aberrant activation of the Wnt signaling pathway in lung cancer. In fact, our *in silico* analysis of 1 926 NSCLC patients indicates a significant increased overall survival associated with high expression levels of *ITF2* and low expression of *DKK1.* The same findings without statistical significance were observed from our “in house” cohorts, probably due to the sample size. However, one of the strengths of our cohort is that it is comprised of fresh frozen samples, enabling us to perform high quality RNA-sequencing in a group of NSCLC patients in order to determine the involvement of the Wnt-pathway components in lung cancer development. The differential expression of *DKK1* within the tumor samples allowed us to perform three different bioinformatics contrasts in order to explore all the possibilities regarding tumor development and the Wnt signaling pathway. Contrast A identified genes with a possible involvement in lung cancer development by comparing differential expression in tumors versus control samples. Contrast B identified alterations in the Wnt pathway in NSCLC tumors and those that could be used as potential therapeutic targets. Finally, contrast C identified genes regulated by the Wnt pathway and others involved in NSCLC development. Using this approach we were able to identify coding genes, non-coding genes and transcripts that had not been functionally characterized previously ^16, 38^. Indeed, in this study we have identified three coding genes, *HOXD9, CLDN6* and *RIOX1*, and one non-coding gene, *XIST*, which could be involved in NSCLC progression through the Wnt signaling pathway. Our data was validated by two alternative methodologies (quantitative real time and NGS), both showing strong positive correlations. From these candidates, we observed a negative correlation of *HOXD9* and *ITF2* expression levels. *HOXD9* was significantly downregulated in tumors with high expression levels of DKK1 and upregulated in tumors compared to controls, indicating that it could be involved in tumor progression through an aberrant activation of the Wnt signaling pathway. We also observed that the tumor samples had higher expression levels of *HOXD9* than the controls, as it has been previously reported ^29^. In addition, patients with lower levels of HOXD9 had better overall survival than those with upregulated expression of this gene. These results are consistent with previous studies linking a high expression of *HOXD9* with glioblastoma and hepatocarcinoma ^24, 36^.

Additional functional analysis showed that *ITF2* overexpression in lung cancer cells H23R decreased the expression of *HOXD9*. Taking into account the basal activation of Wnt pathway in this cell line, we believe that an alternative regulatory mechanism affected by *ITF2* is modulating the expression of *HOXD9*. Reinforcing this hypothesis HOXD9 expression has been shown to also be regulated by epigenetic mechanisms such as the long non-coding RNA *HOTAIR* ^30, 39^, which has been linked to cisplatin-resistance ^23^.

In essence, we have identified *ITF2* as a frequently downregulated gene in cisplatin-resistant cancer cells as well as in NSCLC and ovarian cancer patients. We have also observed a statistically significant relationship with a better response to platinum treatment not only in lung, but also in other epithelial tumors, suggesting that *ITF2* could be used a general epithelial tumor platinum-predictive marker. Moreover, we have defined its implication as a molecular mechanism behind the development of cisplatin resistance in cancer cells probably through the activation of the Wnt-signaling pathway and, several of its downstream effectors genes, providing novel insights into the molecular biology and the cellular mechanisms involved in the acquired resistance to the most widely-used chemotherapy agent, cisplatin. Finally, we have suggested two potential therapeutic targets for further study, *ITF2* and *HOXD9.*

## MATERIALS AND METHODS

### Cell culture and cell-viability assays

The NSCLC and ovarian cancer cell lines H23, H460, OVCAR3 and A2780 were purchased from the ATCC (Manassas, Virginia, USA) and ECACC (Sigma-Aldrich, Madrid, Spain) and cultured as recommended. Their CDDP-resistant variants H23R, H460R, OVCAR3R and A2780R were previously established in our laboratory ^13, 37^. Cisplatin (Farma Ferrer, Barcelona, Spain) was used for CDDP-viability assays. Cells were seeded in 24-well dishes at 40,000 cells/well, treated with increasing doses of CDDP (0, 0.5, 1, 1.5, 2 and 3µg/ml) for an additional 72 or 48 hours and stained as described ^5^. Cell viability comparing sensitive vs. resistant cell lines was estimated relative to the density recorded over the same experimental group without drug exposure at same period of time. Cell authentication is included in Table S5.

### Clinical sample and data collection

We selected a representative number of fresh frozen surgical specimens from University Hospital La Paz (HULP)-Hospital Biobank, totaling 25 NSCLC and 10 ovarian cancer samples, belonging to previously reported cohorts of patients ^37^. Ten adjacent normal tissue (ATT) from NSCLC patients, two additional lung tissue samples of non-neoplastic origin from autopsies and ten more normal ovarian samples obtained from sex reassignment surgery or tubal ligation were used as negative controls (NC). All tumor patients had both a perioperative PET-CT scan showing localized disease and a pathological confirmation of stages after having undergone a complete resection for a histologically confirmed tumor. The samples were processed following the standard operating procedures with the appropriate approval of the Human Research Ethics Committee at IdiPAZ, including informed consent within the context of research. Clinical follow-up was conducted according to the criteria of the medical oncology division, pathological and therapeutic data were recorded by an independent observer and a blind statistical analysis was performed on these data.

### DNA extraction and array of Comparative Genome Hybridization

DNA from cell lines was isolated as previously described ^12^ and used to analyze copy number variations by the Array-CGH SurePrint G3 Human CGH Microarray 1×1M (Agilent, Santa Clara, California, USA). Array experiments were performed as recommended by the manufacturer, described in detail in the GEO repository number GSE129692. The Aberration Detection Method 2 (ADM-2) quality weighted interval score algorithm identifies aberrant intervals in samples that have consistently high or low log ratios based on their statistical score. The score represents the deviation of the weighted average of the normalized log ratios from its expected value of zero calculated with Derivative Log2 Ratio Standard Deviation algorithm. A Fuzzy Zero algorithm is applied to incorporate quality information about each probe measurement. Our threshold settings for the CGH analytics software to make a positive call were 6.0 for sensitivity, 0.45 for minimum absolute average log ratio per region, and 5 consecutive probes with the same polarity were required for the minimum number of probes per region.

### RNA extraction, RT-PCR, qRT-PCR

Total RNA from human cancer cell lines and surgical samples was isolated, reverse transcribed and quantitative RT-PCR analysis was performed as previously described ^37^. For RT-PCR, 2 µl of the RT product (diluted 1:5) was used for semi-quantitative PCR or qPCR reactions with Promega PCR Mix (Promega, Madison, Wisconsin, USA) and SYBR Green PCR Mix (Applied Biosystems, Waltham, Massachusetts, USA), respectively. RT-PCR was performed under the following conditions: (a) 1 cycle of 95°C for 2 min; (b) 30–40 cycles of 95°C for 1 min, 56°C–60°C for 1 min, 72°C for 1 min; (c) an extension of 5 min at 72°C. qRT-PCR absolute quantification was calculated according to the 2^−ΔCt^ method using GAPDH as endogenous control, whereas relative quantification was calculated with the 2^−ΔΔCt^ using *GAPDH* as endogenous control and the sensitive-parental cell line as a calibrator. Samples were analyzed in triplicate using the HT7900 Real-Time PCR system (Applied Biosystems, USA). Primers and probes for qRT-PCR expression analysis were purchased from Applied Biosystems (*TCF4*: Hs00162613_m1; *LRP1B*: Hs01069120_m1; *DKK1*: Hs00183740_m1; *GADPH*: Hs03929097_g1). Primers for *DKK1, HOXD9, CLDN6, XIST* and *RIOX1* for RT-PCR assays were designed, when possible, to analyze the specific transcript that showed significant changes in the RNA-seq; all primers and specific amplification conditions are listed in Table S6.

### NGS (RNA-seq) and Wnt signaling pathways analysis

Total RNA from nine tumor tissues, three lung adjacent normal tissues (ATT) from NSCLC samples and two tissue samples of non-neoplastic origin from autopsies were sent to Sistemas Genómicos Company (Valencia, Spain) for RNA-sequencing. Library samples were prepared and sequenced as recommended by the manufacturer (Illumina, San Diego, California, USA) described in detail in the GEO repository number GSE127559. The bioinformatics analysis was performed in the HULP. Reads were analyzed to quantify genes and isoforms through the RSEM-v1.2.3 methodology (RNA-seq by Expectation Maximization) ^21^ and using the hg19 versions as reference for annotation. The differential expression was carried out with edgeR, which estimates the common and individual dispersion (CMN and TGW, respectively) to obtain the variability of the data ^31^. P-values and FDR statistical analysis were performed by Cmn and Twg models and statistical cut-off point was set as FDR<0.05. Normalization was performed by the TMM method (Trimmed mean of M-values) ^32^. The bioinformatics analysis included an efficiency analysis for every sample, considering the total efficiency as the percentage of reads annotated belonging to a transcript regarding the total fragments initially read. When using Principal Component Analysis (PCA), no differences were observed between samples from non-neoplastic autopsies and adjacent normal tissue (ATT) from NSCLC patients in terms of transcriptomic profile, therefore both types of samples were considered as reference groups for the differential expression analysis (Figure S5). Three different bioinformatics contrasts are described in detail in the results section.

### Transfection assays: top-fop, B-cat and TCF4 Overexpression

A Myc-DDK-tagged ORF clone of *TCF4* and the negative control pCMV6 were used for *in transient* transfection (RC224345; OriGene, Rockville, Maryland, USA) using previously described methodology ^37^. Cells were plated onto 60-mm dishes at 600 000 cells/dish and transfected with a negative control or *TCF4* vectors, using jet-PEI DNA Transfection Reagent (PolyPlus Transfection, Illkirch, France). For the Wnt reporter assay cells were plated at a concentration of 600 000 cells/well in 6-MW plates. Cells were serum starved overnight and co-transfected with 0.2 µg of either Super8xTopFlash (containing 7 copies of the TCF/LEF binding site) or Super8xFopFlash (containing 6 mutates copies of the TCF/LEF binding sites) expression plasmids, and 0.1µg pRL-TK (Renilla-TK-luciferase vector, Promega, USA) as a control, using Lipofectamine 2000. Cells were subsequently treated with LiCl 10mM or co-transfected with S33Y B-catenin for 48 hours prior to luciferase activities being measured using a Glomax 96 Microplate Luminometer (Turner Biosystems Instrument, Sunnyvale, California, USA). Firefly luciferase activity was calculated as light units normalized with the Renilla activity generated by the pRL-CMV vector. B-catenin/TCF activity was calculated by obtaining the ratio of the Top/Fop promoter activities and expressed in relative terms as the fold change of the untreated cells activation levels (=1).

### In silico databases: The Cancer Genome Atlas and Kaplan Meier Plotter

#### The Cancer Genome Atlas (TCGA) data

We selected samples from the TCGA data sets containing RNA sequencing information: 230 lung Adenocarcinoma, 487 lung SCC, 186 Esophageal adenocarcinoma and 522 Head and Neck SCC tumors, to analyze the expression of *ITF2* and *HOXD9* genes. We obtained overall survival Kaplan-Meier estimations selecting groups of patients with high and low expression of *ITF2/HOXD9* (mRNA expression Z-score threshold=1). Samples with *ITF2* deletion and low expression were included in the *ITF2* low group.

#### Kaplan-Meier Plotter

We obtained the survival analysis of 1,926 patients from the Kaplan-Meier Plotter online tool^9^ for the affymetrix ID probes 213891_s_at for *ITF2*, 204602_at for *DKK1* and 206604_at for *HOXD9.* Groups were separated according to the median in high and low expression.

### Statistical analysis

Data were compared using the chi-squared test or Fisher’s exact test for qualitative variables, and Student’s T-test or the Wilcoxon-Mann-Whitney test for quantitative variables. Correlation of quantitative variables was analyzed by Pearson’s test. Overall survival was estimated according to the Kaplan-Meier method and compared between groups by means of the Log Rank test. All the p-values were two-sided, and the type I error was set at 5 percent. Statistical analyses were performed using SPSS_20 software (IBM, Armonk, New York, USA).

## ACKNOWLEDGEMENTS

The authors thank Hayley Pickett for the English language correction. The authors also acknowledge HULP Biobank for sample processing.

## AUTHOR CONTRIBUTIONS

IIC: Conception design and draft

OP, ASP, AGG, MPB, CR, RMA, VP and OVP: development of methodology

OP, ASP, CRA, AGG, RR, DSC, CR, JdC and IIC: acquisition of data

OP, ASP, AGG, RR, DSC, CR, PS, JdC, OVP and IIC: analysis and interpretation of data

CRA and MPB: bioinformatics analysis and interpretation of bioinformatics data OVP: wrote the manuscript

All authors reviewed and/or revised the manuscript.

IIC and OVP: Approved the final version

IIC and OVP: Agreed to be accountable for all aspects of the work in ensuring that questions related to the accuracy or integrity of any part of the work are appropriately investigated and resolved.

