## Supplementary Figures and Tables for "A novel role for the tumor suppressor gene *ITF2* in lung tumorigenesis and chemotherapy response"

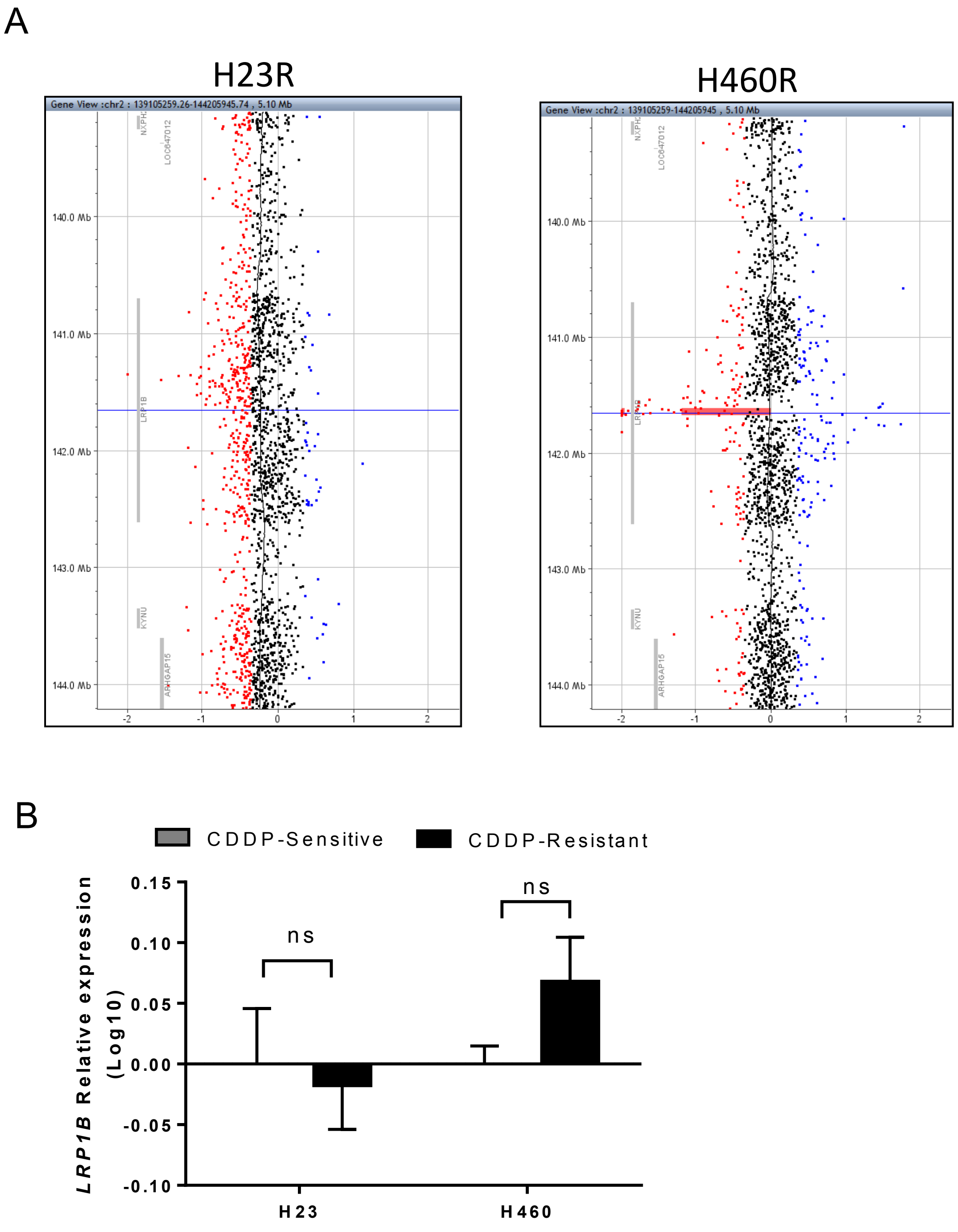


**Supplementary Figure 1: *LRP1B* deletion and expression levels in H23 and H460 cell lines.** (A) Picture extracted from the Agilent Cytogenomics 3.0.1.1 software showing the *LRP1B* deletion in chromosome 2 in H23R and H460R cell lines. (B) Relative mRNA expression levels of *LRP1B* measured by qRT-PCR. The results show the mean fold induction compared to the sensitive cells. Gene expression was normalized to *GAPDH*. Data represent the relative expression levels obtained from the combination of two independent experiments measured in triplicate ± SD. ns: not significant.

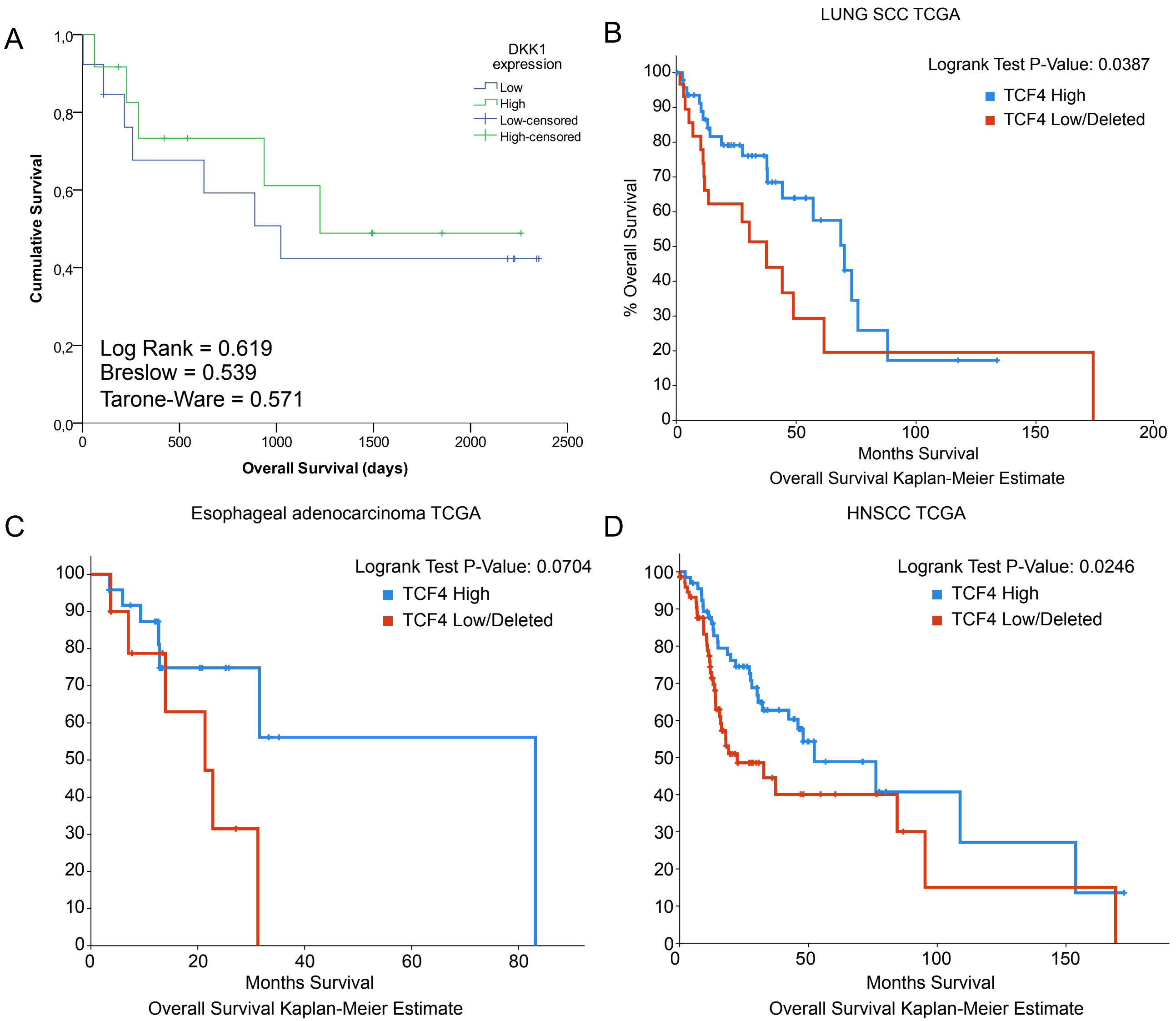


**Supplementary Figure 2:** (A) Kaplan-Meier analysis in a cohort of 25 NSCLC patients from Hospital la Paz for *DKK1* expression. LogRank, Breslow and Tarone-Ware tests were used for comparisons and $p<0.05$ was considered as a significant change in OS. (B-D) Kaplan-Meier analysis on TCGA datasets from lung SCC (n=230; High n=48; Low/Del n=33)(B), esophageal adenocarcinoma (n=186: High n=25; Low/Del n=41)(C) and head and neck SCC (n=522; High n=66; Low/Del n=74)(D) comparing tumors with deletion or low expression of *TCF4* compared with tumor with increased *TCF4* expression.

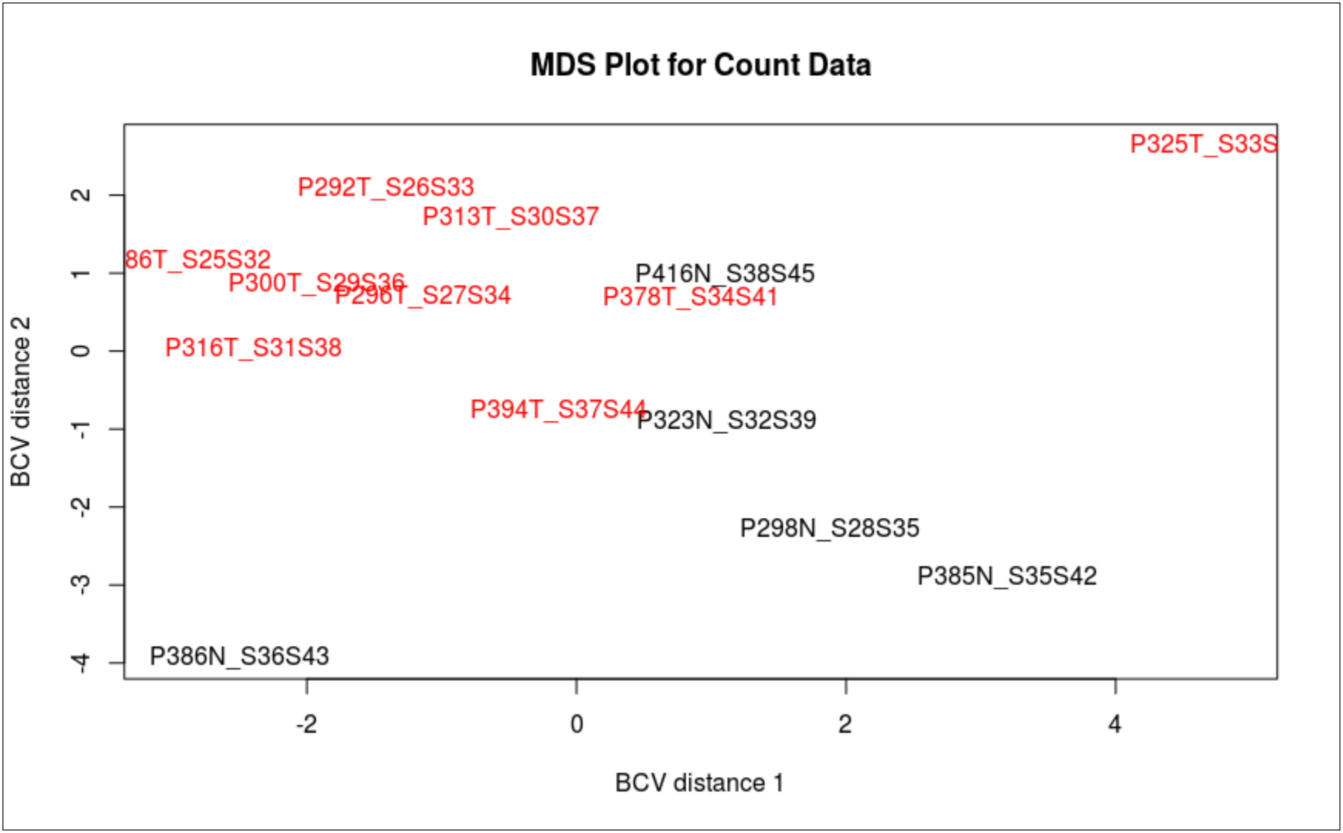


**Supplementary Figure 5. Plot samples on a two-dimensional scatterplot.** In this plot, distances are based on biological coefficient of variation (BCV, square root of the common dispersion). A set of the 500 most differentially expressed genes with the largest biological variation among the libraries were chosen for the analysis. Samples P386 and P385 are lung tissue samples of non-neoplastic origin from autopsies. Samples P298, P323 and P416 are adjacent normal tissue (ATT) from NSCLC patients.

Supp.Figure4

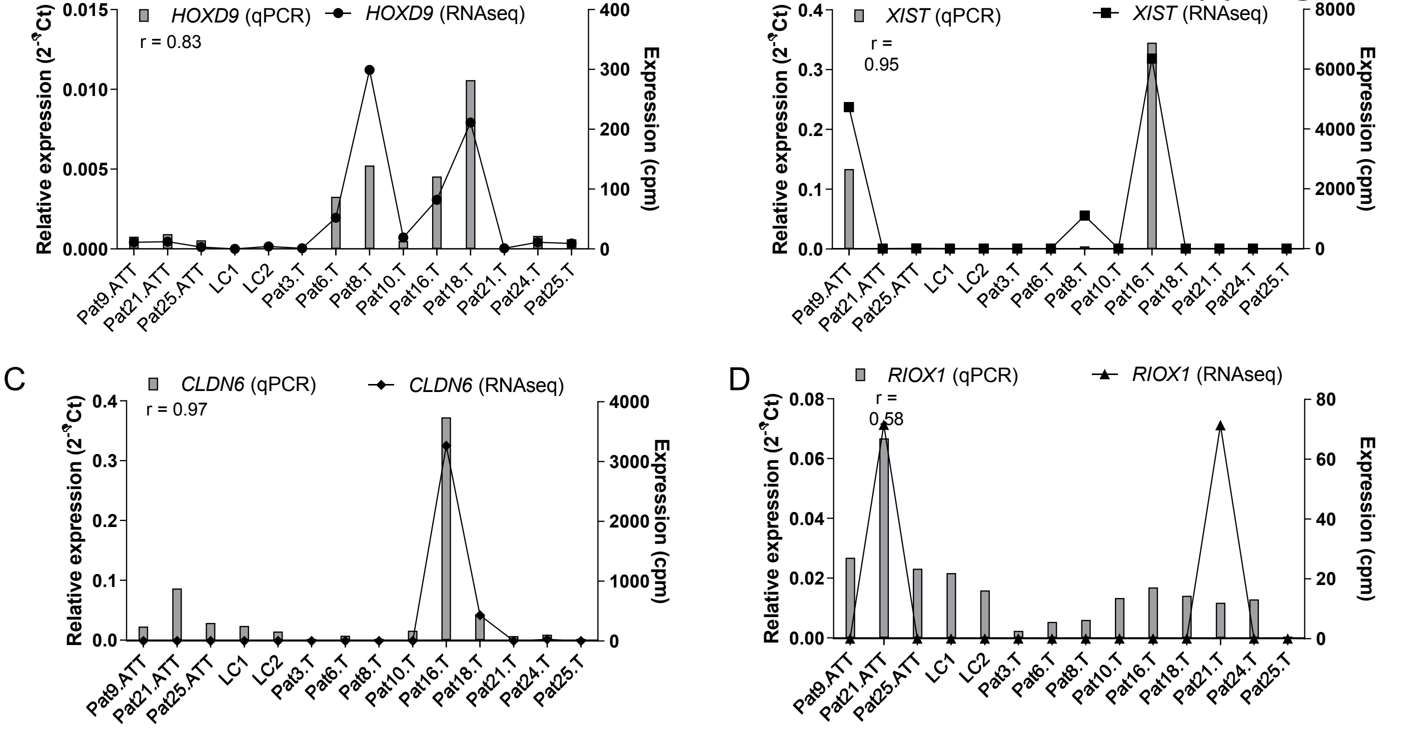


**Supplementary Figure 4. Expression levels of candidate target genes measured by RNA-seq and qRT-PCR in tumor and non-tumor samples from NSCLC patients.** (A-D) correlation between RNA-seq and qRT-PCR expression levels performed in a selection of 14 samples in *HOXD9* (A), *XIST* (B), *CLDN6* (C) and *RIOX1* (D). Bars represent the relative expression of each gene measured by qRT-PCR in triplicate and represented as $2^{-\Delta Ct}$ and lines represent the count per million obtained from the RNA-seq analysis.

**Supplementary Table 1. Summary of the common deleted regions and genes in resistant cells. bp (base pairs)**

| Chromosome | Cytoband | H23/H460<br>Deleted region | A2780/OVCAR3<br>Deleted region | H23/A2780/OVCAR3<br>Deleted region | Common Nº bp | Common Genes |
|---|---|---|---|---|---|---|
| chr2 | q22.1 | 141618089-141673466 | | | 55377 | *LRP1B* |
| chr9 | q22.33 | | 99315964-99362235 | | 46271 | *TMOD1* |
| chr18 | q21.2 - q21.31 | | | 50304876-53014765 | 2709889 | *C18orf26, RAB27B, CCDC68, TCF4, TXNL1, WDR7, BOD1P* |
| chr18 | q21.32 | | | 54661447-55069272 | 407825 | *ZNF532, LOC390858, SEC11C, GRP* |

**Supplementary Table 2.** Clinicopathological and experimental data obtained from patients with ovarian cancer from La Paz University Hospital

| Patient | Histology | Stage | Grade | Chemotherapy | Status | OS (days) | *ITF2* ($2^{-\Delta\Delta Ct}$) | *DKK1* ($2^{-\Delta\Delta Ct}$) |
|---|---|---|---|---|---|---|---|---|
| 1T | Serous | IIIC | Differentiated | CBDCA+Paclitaxel | Alive | 813 | 0.102 | 2.32 |
| 2T | Serous | NA | Differentiated | No | Exitus | 1060 | 0.113 | 0.41 |
| 3T | Serous | NA | Poorly differentiated | No | Exitus | 14 | 0.045 | 0.033 |
| 4T | Serous | IIC | Poorly differentiated | CBDCA+Paclitaxel | Alive | 1632 | 0.02 | 0.041 |
| 5T | Endometrioid | IA | Differentiated | No | Alive | 1736 | 0.048 | 0.09 |
| 6T | Serous | III | Poorly differentiated | CDDP+Paclitaxel | Alive | 1013 | 0.023 | 0.056 |
| 7T | Serous | IV | Poorly differentiated | CBDCA+Paclitaxel | Exitus | 498 | 0.041 | 0.045 |
| 8T | Clear Cell | III | NA | CBDCA+Paclitaxel | Exitus | 1135 | 0.059 | 0.231 |
| 9T | Serous | IIIC | Poorly differentiated | CBDCA+Paclitaxel | Exitus | 716 | 0.122 | 0.354 |

*Note: OS, Overall Survival; CDDP: cisplatin; CBDCA: carboplatin; NA: not available.*

**Supplementary Table 3.** Clinicopathological and experimental data of *ITF2, DKK1, CDLN6* and *XIST* for the NSLC patients that were used for RNAseq

| Patient | Histology | Sex | Stage | Chemotherapy | Relapse | Status | OS (days) | PFS (days) | *ITF2* ($2^{-\Delta Ct}$) | *DKK1* ($2^{-\Delta Ct}$) | *CLND6* ($2^{-\Delta Ct}$) | *XIST* ($2^{-\Delta Ct}$) | *RIOX1* ($2^{-\Delta Ct}$) |
|---|---|---|---|---|---|---|---|---|---|---|---|---|---|
| Pat3.T | Epidermoid | Male | IB | No | Yes | Exitus | 1022 | 825 | 0.27 | 1.25 | 1.18·10-3 | 2.71·10-7 | 2.33·10-3 |
| Pat6.T | Large cell | Male | IIB | No | No | Exitus | 62 | 62 | 0.43 | 1.97 | 7.43·10-3 | 3.92·10-6 | 5.39·10-3 |
| Pat8.T | Epidermoid | Female | IIIB | CDDP + Others | No | Exitus | 109 | 109 | 0.18 | 0.43 | 3.63·10-4 | 4.16·10-3 | 6.03·10-3 |
| Pat10.T | Epidermoid | Male | IB | No | No | Alive | 1853 | 1853 | 0.42 | 2.78 | 1.58·10-2 | 7.09·10-3 | 1.43·10-2 |
| Pat16.T | Adenocarcinoma | Female | IIIA | CDDP + Others | ND | Alive | 2228 | 2228 | 0.24 | 0.08 | 3.73·10-1 | 3.45·10-1 | 1.69·10-2 |
| Pat18.T | Epidermoid | Male | IIB | CBDCA + Others | No | Exitus | 259 | 259 | 0.15 | 0.03 | 4.31·10-2 | 2.67·10-4 | 1.41·10-2 |
| Pat21.T | Adenocarcinoma | Male | IIIA | CDDP + Others | No | ND | 421 | 421 | 0.20 | 4.72 | 6.92·10-3 | 4.32·10-4 | 1.18·10-2 |
| Pat24.T | Adenocarcinoma | Male | IIA | CBDCA + Others | No | Alive | 1491 | 1491 | 0.98 | 10.34 | 9.09·10-3 | 3.64·10-4 | 1.29·10-2 |
| Pat25.T | Adenocarcinoma | Male | IIA | CDDP + Others | No | Alive | 1496 | 1496 | 0.77 | 137.16 | 5.91·10-7 | 1.93·10-5 | 1.70·10-5 |

**Supplementary Table 4.** Selected genes potentially involved in the Wnt signaling pathway identified through a global transcriptomic analysis on 14 samples of NSCLC patients

| **Candidates** | **Contrast A** | | **Contrast B** | | **Contrast C** | | **Statistical Significance** 1=FDR<0.05; 0=FDR>0.05 | | |
|---|---|---|---|---|---|---|---|---|---|
| Gene ID | logFC | FDR | logFC | FDR | logFC | FDR | A | B | C |
| ***HOXB8*** | 4.529 | 0.047 | -4.644 | 0.035 | 1.352 | 0.547 | 1 | 1 | 0 |
| ***HOXD9*** | 3.615 | 0.042 | -3.723 | 0.027 | 1.317 | 0.518 | 1 | 1 | 0 |
| ***CLDN6*** | 8.822 | 0.033 | -7.720 | 0.022 | 2.724 | 0.309 | 1 | 1 | 0 |
| ***AC022596.6*** | 7.159 | 0.046 | -7.566 | 0.028 | 1.207 | 0.628 | 1 | 1 | 0 |
| ***SLC13A2*** | 0.921 | 0.833 | -6.648 | 0.004 | -4.199 | 0.024 | 0 | 1 | 1 |
| ***XIST*** | -0.171 | 0.943 | -11.413 | 4.39E-08 | -10.025 | 0.003 | 0 | 1 | 1 |
| ***RIOX1*** | -0.785 | 0.843 | -7.632 | 0.016 | -6.807 | 0.026 | 0 | 1 | 1 |
| ***AL591684.1*** | -0.022 | 0.971 | -7.147 | 0.022 | -5.554 | 0.018 | 0 | 1 | 1 |
| ***PTPN20A*** | -0.643 | 0.751 | -3.996 | 0.043 | -3.162 | 0.005 | 0 | 1 | 1 |

Note: logFC: Log of FoldChange, FDR: False Discovery Rate.

### Supplementary Table 5. Cell Authentication. Genomics Core Facility. IIBm CSIC-UAM.

| Name of Cell line | Cancer type | Testing Date | | | | | | | | | | | | REF | Match/Not Match |
|---|---|---|---|---|---|---|---|---|---|---|---|---|---|---|---|
| Sample | | | *M musculus* | D5S818 | D13S317 | D7S820 | D16S539 | VWA | TH01 | AMEL | TPOX | CSF1PO | D21S11 | | |
| REF NCI-H23 | | | Negative | 12,13 | 12 | 9,10 | 11 | 16,17 | 6 | X | 8,9 | 10 | | ATCC ® CRL-5800 | |
| H23 | Lung | 06/22/2016 | ******** | 12,13 | 12 | 9,10 | 11 | 16,17 | 6 | X | 8,9 | 10 | *30* | | Match |
| REF NCI-H460 | | | Negative | 9,10 | 13 | 9,12 | 9 | 17 | 9.3 | X,Y | 8 | 11,12 | | ATCC ® HTB-177 | |
| H460 | Lung | 06/22/2016 | ******** | 9,10 | 13 | 9,12 | 9 | 17 | 9.3 | X,Y | 8 | 11,12 | *30* | | Match |
| REF A2780 | | | Negative | 11,12 | 12,13 | 10 | 11,13 | 15,16 | 6 | X | 8,10 | 10,11 | *27,28* | SIGMA | |
| A2780 | Ovary | 06/22/2016 | ******** | 11,12 | 12,13 | 10 | 11,12,13 | 15,16 | 6 | X | 8,10 | 10,11 | *28* | | Match |
| REF OVCAR3 | | | Negative | 11,12 | 12 | 10 | 12 | 17 | 9,9,3 | X | 8 | 11,12 | | ATCC ® HTB-161 | |
| OVCAR3 | Ovary | 06/22/2016 | ******** | 11,12 | 12 | 10 | 12 | 17 | 9,9,3 | X | 8 | 11,12 | *29,31,2* | | Match |

**Method:**

| | |
|---|---|
| STR amplification kit | GenePrintR 10 System (Promega) |
| STR profile analysis software | GeneMapper® v3.7 (LifeTechnologies) |
| Genomic Analyzer System | ABI 3130 XL (Applied Biosystems) |
| DNA source | Cultured cells; cultured cells pellet |
| DNA isolation method | DNeasy blood and tissue kit (Qiagen) |
| DNA quantification method | Qubit 2.0 Fluorometer (Life Technologies) |
| Amount of DNA/amplification | 4 ng |

The GenePrint® 10 System allows co-amplification and three-color detection of ten human loci: TH01, TPOX, vWA, Amelogenin, CSF1PO, D16S539, D7S820, D13S317, D21S11 and D5S818. These loci collectively provide a genetic profile with a random match probability of 1 in 2.92 × 109 and are used for human cell line and tissue authentication and identification and human cell line cross-contamination determination. STRs profiles are sent for comparison with cell line date bases like ATCC (American Type Culture Collection), DSMZ (Deutsche Sammlung von Mikrorganismen and Zellkulturen).

**Supplementary Table 6: List of designed primers used for RT-PCR and the amplification characteristics.**

| GeneSymbol | cDNA Primer Forward | cDNA Primer Reverse | cDNA Amplification Length | Annealing Temperature | Cycles of amplification |
|---|---|---|---|---|---|
| *DKK1* | CAACTACCAGCCGTACCCGTG | AACAGAACCTTCTTGTCCTTTG | 331 | 59ºC | 36 |
| *HOXD9* | AGCAGCAACTTGACCCAAACAACC | TGACCTGTCTCTCTGTTAGGTTGAG | 189 | 59ºC | 36 |
| *CLDN6* | ATCTCCTTCGCAGTGCAGCTCCTTCAAC | TGAGTCGTACACCTTGCACTGCATC | 241 | 59ºC | 36 |
| *XIST* | ATTATAAGTGACCACAGCCATGCAC | TTCATGTCCAAGGTGAGTGCCTATGCTC | 334 | 58ºC | 36 |
| *RIOX1* | ACCGCTGACCTGGATTCGATGCTGCG | AACGTTGGAGCCTGCCATGCTTCC | 252 | 62ºC | 34 |
| *GAPDH* | GAGAGACCCTCACTGCTG | GATGGTACATGACAAGGTGC | 135 | 58ºC | 25 |
